# Corticospinal Tractometry and Whole-Brain Connectometry of Hand Dexterity in Chronic Stroke and Traumatic Brain Injury

**DOI:** 10.64898/2026.08.09.743688

**Authors:** Vikram Shenoy Handiru, Easter S. Suviseshamuthu, Olga Boukrina, Glenn Wylie, Guang H Yue

**Affiliations:** Center for Mobility and Rehabilitation Engineering Research, Kessler Foundation, West Orange, 07052, NJ, USA; Center for Stroke Rehabilitation Research, Kessler Foundation, West Orange, 07052, NJ, USA; Rocco Ortenzio Neuroimaging Center, Kessler Foundation, West Orange, 07052, NJ, USA; Department of Physical Medicine and Rehabilitation, Rutgers New Jersey Medical School, Newark, 07107, NJ, USA

**Author notes:** Contributing authors.

**Keywords:** Diffusion tensor imaging, hand dexterity, connectometry, partial least squares regression, stroke, traumatic brain injury

## Abstract

Hand dexterity impairment is a major contributor to long-term disability after acquired brain injury, yet the white matter substrates supporting residual dexterity remain incompletely understood. We investigated diffusion MRI markers of hand dexterity in individuals with chronic stroke (*n* = 9) and traumatic brain injury (TBI; *n* = 8) using complementary tract-specific and whole-brain approaches. Partial least squares regression (PLSR) was used to evaluate the cross-validated predictive relevance of bilateral corticospinal tract (CST) diffusion and tractometry features, while quantitative anisotropy (QA)-based correlational tractography was used to identify distributed white matter pathways associated with dexterity performance measured using Box and Block Test (BBT) and MusicGlove Dexterity Test(MGDT). In stroke, CST features predicted BBT performance (Q**^2^** = 0.69, r = 0.85, permutation *p* = .010) and, more modestly, MGDT performance (Q**^2^**= 0.22, r = 0.72, permutation *p* = .008). In contrast, CST-based models showed no predictive relevance for dexterity outcomes in TBI. Whole-brain connectometry revealed that better dexterity after stroke was associated with greater QA across distributed pathways extending beyond the CST, including commissural, association, and projection fibers. Box and Block Test performance was prominently associated with callosal and cingulum-related pathways, whereas MusicGlove performance showed greater representation of CST and projection pathways. In TBI, significant connectometry findings for the BBT similarly implicated distributed commissural and association pathways, whereas no significant pathways were identified for the MusicGlove test. Together, these findings suggest that the structural correlates of hand dexterity extend beyond the CST and vary across dexterity measures and injury populations. Although preliminary given the small cohorts, the complementary tractometry and connectometry findings support a network-level characterization of residual hand function after acquired brain injury and motivate validation in larger cohorts.

## 1 Introduction

### Clinical Significance of Hand Dexterity Impairment After ABI

Upper-limb motor impairment remains one of the most persistent and disabling sequelae of acquired brain injury (ABI), including both stroke and traumatic brain injury (TBI). In particular, impaired hand dexterity limits independence in activities of daily living and substantially reduces quality of life [1]. Although many individuals recover basic motor function, about 40–50% continue to exhibit residual impairment six months after onset [2]. Despite its clinical importance, the structural basis of impaired dexterity after ABI remains incompletely understood, especially when comparing focal injuries such as stroke [3] with the more distributed pathology observed in TBI [4, 5].

### Corticospinal Tract Integrity as a Focal Substrate of Motor Function

From a neuroanatomical perspective, the corticospinal tract (CST) is the principal descending pathway mediating voluntary motor control, and its structural integrity is strongly associated with upper-limb function after stroke [6]. Structural or physiological disruption of the CST, particularly within the posterior limb of the internal capsule (PLIC) or cerebral peduncle, is consistently associated with poor upper-limb recovery [7, 8]. Likewise, transcranial magnetic stimulation (TMS) studies demonstrate elevated resting motor thresholds and absent motor-evoked potentials (MEPs) in both stroke and TBI when corticospinal conduction is impaired [9]. Together, these findings identify CST integrity as a key biomarker of residual descending motor drive and motivate a focused analysis of tract-specific diffusion markers related to hand motor function.

### Multivariate CST-specific Diffusion and Tractography Features relevant to Motor Function

Diffusion tensor imaging (DTI) provides quantitative markers of white-matter (WM) microstructure and has been widely used to study motor impairment. Common diffusion metrics such as fractional anisotropy (FA), axial diffusivity (AD), and radial diffusivity (RD) capture complementary aspects of tissue organization, including axonal coherence, fiber architecture, and myelin integrity [10]. Reduced FA in the ipsilesional CST has repeatedly been linked to diminished MEPs and poor motor outcomes [11, 12]. Beyond conventional tensor-derived metrics, higher-order and tractography-derived measures may provide additional information about pathway integrity. Quantitative anisotropy (QA), derived from the spin distribution function, reflects the density of anisotropic diffusion along specific fiber orientations and is less sensitive than FA to crossing fibers and partial-volume effects [13, 14]. The isotropic diffusion component (ISO) may reflect extracellular free water and edema-related signal contributions, thereby providing complementary information about tissue microenvironment not captured by anisotropy metrics alone [15, 16], whereas fiber count provides a tractography-derived estimate of pathway continuity [17]. In addition, asymmetry indices comparing ipsilesional and contralesional CST properties may better capture pathological deviations associated with Wallerian degeneration and motor impairment [18]. Moreover, studies have shown that patients with more pronounced asymmetry may experience greater relative gains under appropriate rehabilitation conditions, indicating a non-linear relationship between baseline tract integrity and motor improvements [19, 20].

Because these CST-related features are numerous and potentially collinear, a multivariate framework is needed to evaluate their joint relationship with behavior. Partial least squares regression (PLSR) is well suited for this purpose because it identifies latent variables that maximize the covariance between imaging features and behavioral outcomes while reducing the impact of collinearity and multiple-comparison inflation [21–23]. In the present study, PLSR was used to examine how tract-specific diffusion features indexing CST integrity relate to manual dexterity and corticospinal excitability.

### Distributed Pathway Organization Beyond the CST

Direct corticospinal projections play a central role in individual finger movements, which are fundamental to fine motor control and dexterity [24]. Consistent with this, experimental and clinical studies demonstrate that damage to the motor cortex or CST selectively impairs the ability to produce independent finger movements and thereby affects hand dexterity, even when more global upper-limb function is relatively preserved [25]. While CST integrity is critical for voluntary motor function, manual dexterity is not solely determined by CST integrity. Dexterous hand function extends beyond isolated motor output and emerges from coordinated activity across a distributed sensorimotor network, including primary motor and somatosensory cortices, premotor and posterior parietal regions, basal ganglia, cerebellum, and the white-matter pathways that support communication among these regions. This network-level organization is particularly relevant for clinical measures of dexterity that involve complex, goal-directed tasks requiring visuomotor integration, somatosensory feedback, motor planning, and interhemispheric coordination.

Accordingly, dexterity impairments following acquired brain injury may reflect disruption of both descending motor pathways and broader structural networks. This broader network perspective is especially relevant in TBI, where white-matter damage is often diffuse and heterogeneous, in contrast to the more focal, regionally localized lesions typically observed in stroke [4, 5]. Converging evidence further suggests that incorporating distributed connectivity measures, including frontoparietal, cortico-reticulospinal, and cerebello-thalamo-cortical pathways, alongside CST-specific features improves the characterization of motor outcomes [8, 26–28]. These observations motivate a complementary, whole-brain analysis to determine whether dexterity is associated with distributed white-matter pathway organization beyond focal CST integrity.

### Diffusion MRI Connectometry as a Complementary Analysis of Distributed Pathways

Diffusion MRI connectometry provides a data-driven approach for identifying white-matter pathways associated with behavioral variability without requiring *a priori* tract selection. Unlike conventional tract-based analyses, which summarize diffusion properties within predefined regions of interest, connectometry evaluates local connectome features along white-matter pathways and tracks segments whose microstructural properties are statistically associated with behavioral measures. In the present study, these associations were quantified using Spearman partial correlation while controlling for age and time since injury, thereby isolating structure–function relationships independent of these known sources of variability. This approach is particularly useful for detecting distributed structure–function relationships in conditions characterized by heterogeneous or network-level disruption. Recent studies have shown that behaviorally relevant information may be distributed across commissural, association, and projection pathways rather than confined to canonical tracts alone. For example, network-level associations between interhemispheric connectivity and behavioral recovery have been reported in stroke [29, 30], whereas broader patterns of structural disconnection have been linked to cognitive deficits in TBI [31, 32]. Thus, connectometry provides a complementary framework for testing whether dexterity is related to distributed WM organization beyond focal CST integrity.

### Study Objective and Hypothesis

The present study investigated diffusion-based predictors of manual dexterity and corticospinal excitability in individuals with chronic stroke and TBI using two complementary analyses. First, a PLSR-based analysis examined whether CST-specific diffusion features—including FA, QA, AD, RD, ISO, fiber count, and CST asymmetry—explain significant variance in physical function and neurophysiological measures relevant to hand motor function. Second, diffusion MRI connectometry was used to identify distributed white-matter pathways whose local structural properties are associated with behavioral variability. We hypothesized that CST-specific diffusion features would significantly relate to dexterity and corticospinal excitability, and that connectometry would reveal additional distributed pathway associations beyond focal CST integrity alone. Overall analysis framework is illustrated in Fig. 1.

**Fig. 1.**
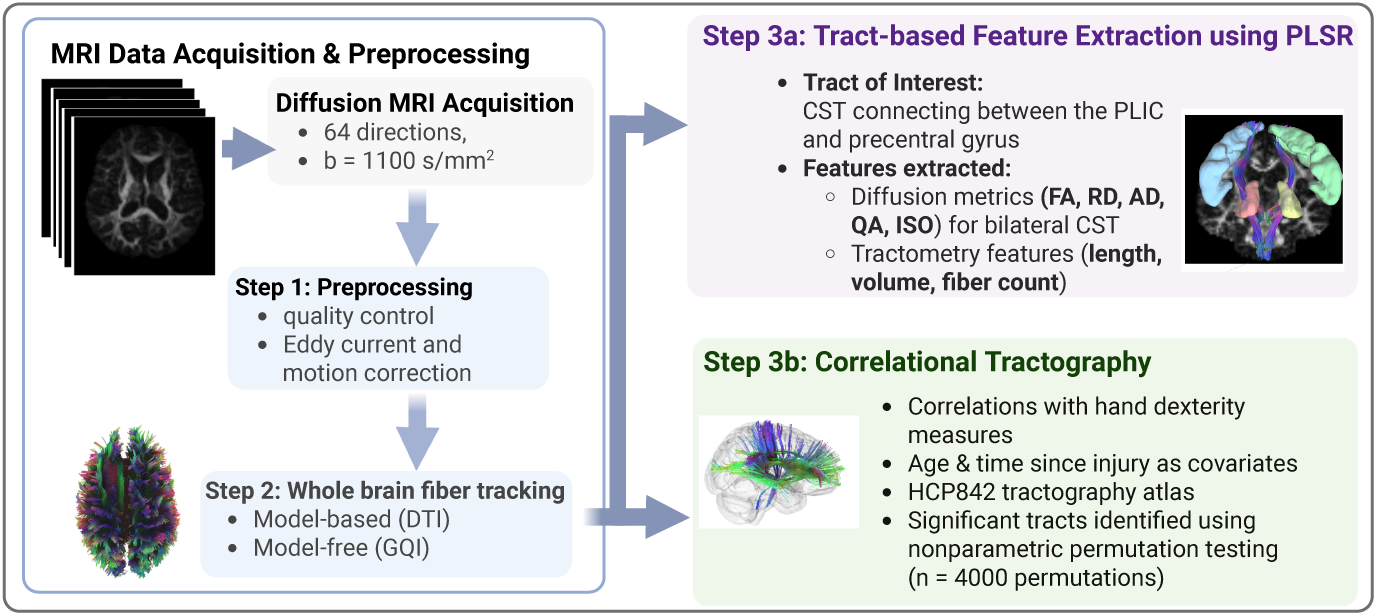
Diffusion MRI analytic workflow for tract-based feature analysis and correlational tractography. Diffusion MRI data were processed using DSI Studio software, which included eddy-current and motion artifacts correction, followed by whole-brain fiber reconstruction using both DTI- and GQI-based approaches. The pipeline then branched into two complementary analyses: (i) tract-based extraction of bilateral corticospinal tract features between the PLIC and precentral gyrus, including diffusion metrics (FA, RD, QA, ISO) and tractometry measures (length, volume, fiber count), which were used to predict hand outcome measures using partial least squares regression (PLSR); and (ii) whole-brain correlational tractography to identify white matter pathways associated with dexterity. Connectometry findings were evaluated using nonparametric permutation testing (4000 permutations) with FDR correction and tract labeling with the HCP842 atlas. Abbreviations: **DTI**: Diffusion Tensor Imaging; **GQI**: Generalized Q-sampling Imaging; **PLIC**: Posterior limb of the internal capsule; **FA**: Fractional Anisotropy; **RD**: Radial Diffusivity; **QA**: Quantitative Anisotropy; **ISO**: Isotropic diffusion

## 2 Methods

### 2.1 Participants

Participants were drawn from two independent studies conducted at Kessler Foundation (New Jersey, USA) during the period 2019-2023. Data collection was interrupted during the COVID-19 pandemic, during which in-person research visits were temporarily paused. These pilot studies investigated the effects of targeted noninvasive brain stimulation methods on the hand motor function after acquired brain injury. For the present manuscript, only baseline data acquired before any intervention or follow-up assessment were analyzed; therefore, the present work should be interpreted as a retrospective, cross-sectional baseline imaging-behavior analysis. All study procedures were approved by the Kessler Foundation Institutional Review Board (Protocols R-1056-19 and R-1067-19), and all participants provided written informed consent before participating. Because the present analysis used deidentified retrospective baseline data, only datasets meeting the required MRI and behavioral data availability criteria were included.

The analytic sample included adults with chronic stroke (n = 9) or chronic TBI (n = 8) who were 18 to 75 years old, at least 6 months post-injury, medically stable, and able to complete the study assessments. Eligible participants had mild-to-moderate hand impairment, operationally defined at screening by a Box and Block Test (BBT) score lower than 50 blocks per minute. This threshold corresponds to performance at least 2 standard deviations below age-matched normative values in healthy adults [33] and is consistent with mild-to-moderate distal motor impairment reported in prior stroke cohorts [34]. Exclusion criteria included severe hand spasticity (Modified Ashworth Scale score *≥* 3), substantial sensory loss in the affected hand, neurological conditions other than stroke or TBI, contraindications to MRI, and contraindications to transcranial magnetic stimulation (TMS), including tinnitus, recurrent seizures or epilepsy, or seizure-threshold-lowering medications. The study size was determined by the parent pilot studies rather than by a formal sample-size calculation for the present retrospective analysis. The parent studies targeted n = 12 per group, consistent with commonly used pilot-study planning heuristics [35]. The final sample size for the present analysis was determined by the availability of complete baseline diffusion MRI, behavioral, and neurophysiological data suitable for the planned analyses.

Because these data were derived retrospectively from two independent small cohorts, the stroke and TBI groups were not matched for demographic or injury-related characteristics. As reported in Table 1, the groups differed in age and time since injury, and the TBI cohort was exclusively male. The stroke lesions were predominantly left hemispheric, corresponding to clinically evident right-hand weakness in the stroke cohort. TBI was not characterized by a single focal lesional hemisphere. Accordingly, all imaging–behavior analyses were conducted separately by cohort.

**Table 1.** Participant demographic, clinical, behavioral, and neurophysiological characteristics.

| Characteristic | Stroke<br>(n=9) | TBI (n=8) | *p*-value |
| --- | --- | --- | --- |
| *Demographic and injury characteristics* |  |  |  |
| Age, years | 57.8 ± 7.2 | 46.8 ± 11.2 | **0.027** |
| Female, n (%) | 4 (44%) | 0 (0%) | 0.082 |
| Male, n (%) | 5 (56%) | 8 (100%) |  |
| Right-handed, n (%) | 6 (67%) | 5 (56%) | 1.000 |
| Time since injury, months | 25.0 ± 12.4 | 186.8 ± 117.8 | **0.003** |
| *Behavioral and neurophysiological measures* |  |  |  |
| BBT, blocks/min | 27.3 ± 17.5 | 33.6 ± 15.1 | 0.432 |
| MGDT, score | 35.4 ± 18.6 | 56.9 ± 14.2 | **0.020** |
| 9HPT, time in s | 74.1 ± 77.8 | 42.3 ± 13.1 | 0.260 |
| FDI MVC, mV | 0.25 ± 0.13 | 0.38 ± 0.23 | 0.174 |
| MEP amplitude, mV | 0.22 ± 0.13 | 0.21 ± 0.16 | 0.950 |
| TMS motor threshold, %MSO | 45.1 ± 20.3 | 50.9 ± 17.4 | 0.552 |

Values are reported as mean ± standard deviation or number (percentage). Reported *p*-values are unadjusted and are provided for describing the cohort characteristics. Age, time since injury, behavioral outcomes, and neurophysiological measures were compared using Welch’s two-sample *t*-tests because of small sample size and unequal variances across groups. Sex and handedness distributions were compared using Fisher’s exact tests because of sparse contingency-table counts. Although the sex distribution comparison did not reach $p < 0.05$ , the TBI cohort consisted exclusively of male participants.

### 2.2 Motor and Neurophysiological Assessments

Baseline clinical assessments consisted of complementary measures of hand function, capturing distinct but interrelated components of hand dexterity. The BBT quantifies rapid unilateral object transfer and was used as a measure of gross manual dexterity and functional hand use [36]. The MGDT was included as an instrumented task-specific measure of finger-level movement timing, coordination, and sequencing during MusicGlove interactions [37]. Although MGDT is not a standardized clinical outcome, it was included because its timing and multi-finger sequencing demands are closely aligned with finger independence and temporal coordination, which are frequently impaired after stroke [38]. The 9HPT was included as a measure of fine dexterity, and it was analyzed as raw completion time in seconds, where longer time indicates worse fine motor performance. First dorsal interosseous (FDI) maximum voluntary contraction (MVC) and TMS-evoked motor evoked potential (MEP) amplitude were treated as secondary exploratory outcomes of neurophysiology relevant to finger movement. BBT and 9HPT were administered bilaterally, with each hand tested separately. For all analyses, only performance from the affected upper limb was used, defined as the limb contralateral to the lesion in the stroke cohort. Because TBI was not characterized by a single focal lesional hemisphere, the affected hand was defined as the clinically more impaired hand determined using BBT scores.

Neurophysiological integrity of the corticospinal system was assessed using MEPs elicited via TMS. MEP amplitude serves as an index of corticospinal excitability and functional integrity, which are critical for voluntary motor output and dexterous hand function. Given that fine motor control relies heavily on intact corticospinal projections and precise neural drive to intrinsic hand muscles, MEP measures provide an important neurophysiological correlate of behavioral dexterity outcomes [39]. MEPs were recorded from the FDI muscle of the affected hand using surface electromyography (PowerLab 8/35, ADInstruments, Australia). To evoke MEPs, single-pulse TMS was delivered using a Magstim D110 stimulator (Magstim, UK) connected to a figure-of-eight coil (D70^2^), with the individual MRI-guided neuronavigation performed using the Brainsight system (Rogue Research, Montreal, Canada) to ensure precise targeting of the primary motor cortex hand representation. Resting motor threshold (RMT) was defined as the minimal stimulator output required to elicit MEPs *≥* 50*µ*V in atleast 6 of 10 trials, and MEP amplitude was averaged across 20 stimuli delivered at 120% RMT [40].

### 2.3 MRI Acquisition and DTI Processing

All neuroimaging procedures were performed using a Siemens Skyra 3T MRI scanner at the Rocco Ortenzio Neuroimaging Center at Kessler Foundation. Structural imaging was acquired using a T1-weighted magnetization-prepared rapid gradient-echo (MPRAGE) sequence (TR = 2.1 s; TE = 3.43 ms; flip angle = 9*^◦^*; field of view = 256 *×* 256 mm^2^; 1-mm isotropic voxels). Diffusion-weighted images were acquired using a two-dimensional single-shot echo-planar imaging sequence with diffusion encoding along 64 non-collinear directions at *b* = 1100 s/mm^2^, together with one non-diffusion-weighted (*b* = 0 s/mm^2^) volume. Additional acquisition parameters were TR = 4600 ms, TE = 81 ms, flip angle = 90*^◦^*, 2-mm isotropic voxels, no interslice gap, and 66 axial slices. Identical diffusion acquisition parameters were used across participants.

Diffusion data were processed in DSI Studio (Chen version). Eddy-current and motion correction were performed through the DSI Studio interface using FSL *eddy*, and b-table orientation was checked against a population-averaged template. Diffusion data were reconstructed using generalized q-sampling imaging with a diffusion sampling length ratio of 1.25. Tensor-derived metrics were calculated using diffusion-weighted volumes with *b <* 1750 s/mm^2^.

Tractography focused on the bilateral corticospinal tracts (CSTs), with the posterior limb of the internal capsule (PLIC) and precentral gyrus (PrG) used as anatomical inclusion regions. Atlas-based ROIs were defined in template space. PLIC masks were derived from the 1-mm Johns Hopkins University International Consortium for Brain Mapping (JHU-ICBM) white-matter labels atlas (https://identifiers.org/neurovault.image:1401), whereas PrG masks were derived from the Automated Anatomical Labeling (AAL) atlas. Following registration, ROI alignment was visually inspected for each participant. When visible misregistration was identified, limited participant-specific adjustment of ROI position in the anterior–posterior direction was performed to restore anatomical correspondence with the PLIC; otherwise, the registered atlas ROIs were retained without modification. No additional manual reshaping of the ROIs was performed.

CSTs were reconstructed using deterministic fiber tracking [14] with augmented tracking strategies. The PLIC and PrG were specified as inclusion ROIs, such that retained streamlines were required to pass through both regions; and no exclusion ROIs were applied. The anisotropy threshold and tracking step size were randomly selected by DSI Studio, and the angular threshold was randomly selected within 15*^◦^*– 90*^◦^*. Tracking was terminated after 1,000,000 seeds. Streamlines shorter than 10 mm or longer than 200 mm were discarded, and topology-informed pruning was applied with two iterations to reduce anatomically implausible or noisy streamlines [41]. Quantitative diffusion indices along reconstructed streamlines were sampled using trilinear interpolation.

For each CST, mean FA, QA, AD, RD, ISO, fiber count, tract length, and tract volume were extracted. Fiber count represents the number of reconstructed streamlines satisfying the tract-of-interest and filtering criteria. Fiber length was calculated as the mean physical length (mm) of the retained streamlines, whereas tract volume represented the total volume of voxels traversed by the reconstructed streamlines. Accordingly, fiber count, tract length, and tract volume were interpreted as reconstruction-derived measures of pathway organization and spatial extent rather than direct biological measures of axonal number, tract strength, or absolute anatomical connectivity.

For the stroke cohort, CST asymmetry was quantified using an asymmetry index (AI). FA asymmetry (*FA_AI_ ∈* [*−*1, 1]) was calculated as:

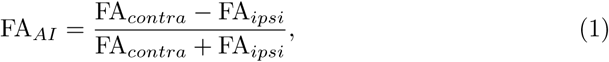

where FA*_ipsi_* and FA*_contra_* denote values from the ipsilesional and contralesional CSTs, respectively, such that positive values indicate lower FA in the ipsilesional CST [42]. Similar metrics were also derived for the QA (*QA_AI_*) and fiber count (*FiberCount_AI_*). Because the TBI cohort did not have a consistently defined lesional hemisphere, ipsilesional and contralesional terminology was not applied when interpreting asymmetry measures in this cohort.

### 2.4 Lesion Analysis in Stroke

To provide anatomical context for the stroke-specific tractography findings, we quantified both lesion topography and lesion-induced structural disconnectivity in the stroke cohort. This analysis was motivated by lesion network mapping approaches showing that focal stroke lesions may disrupt WM connections beyond the lesion area, thereby producing behavioral effects that are not fully captured by lesion location alone [43]. These analyses were restricted to participants with stroke because the TBI cohort did not have a comparable focal lesion phenotype suitable for lesion-overlap or lesion-disconnectivity mapping. Stroke lesion masks were generated from T1-weighted MRI using a pretrained nnU-Net framework developed for chronic stroke lesion segmentation [44–46]. To reduce false-positive predictions outside the intracranial space, SynthStrip-based brain masking was applied as a postprocessing step [47]. Final binary masks were visually inspected by the co-author (Dr. Boukrina) experienced in stroke lesion segmentation and were spatially normalized to MNI152 standard space using affine registration of each participant’s T1-weighted image to the MNI template, followed by nearest-neighbor transformation of the lesion mask.

Lesion-induced disconnectivity was estimated using the Lesion Quantification Toolkit (LQT), a MATLAB-based toolbox, following a lesion-connectome approach similar to prior work in stroke motor recovery [48, 49]. For each participant, the MNI-normalized lesion mask was projected onto the HCP-1065 population tractography atlas to estimate the proportion of atlas streamlines intersecting lesioned tissue. This procedure generated subject-level voxel-wise tract disconnection maps, which were subsequently averaged to visualize the group-level burden of lesion-induced WM disconnection. These lesion and disconnectivity maps were only used descriptively to support anatomical interpretation of the PLSR and connectometry findings.

### 2.5 Data Analysis

#### 2.5.1 Partial Least Squares Regression Modeling

Tract-level diffusion and tractometry features were used to predict hand function outcomes with PLSR, as illustrated in Fig. 2. Predictors included bilateral CST-derived FA, QA, AD, RD, ISO, fiber count, tract length, tract volume, and asymmetry indices. BBT and MGDT were designated as the primary behavioral dexterity outcomes; 9HPT, FDI MVC, and MEP amplitude were secondary exploratory outcomes. Outcome-specific complete-case analysis was used, and no missing values were imputed. PLSR was selected because the diffusion predictors were expected to be collinear and because the sample size was too small to support conventional multivariable regression with many correlated predictors.

**Fig. 2.**
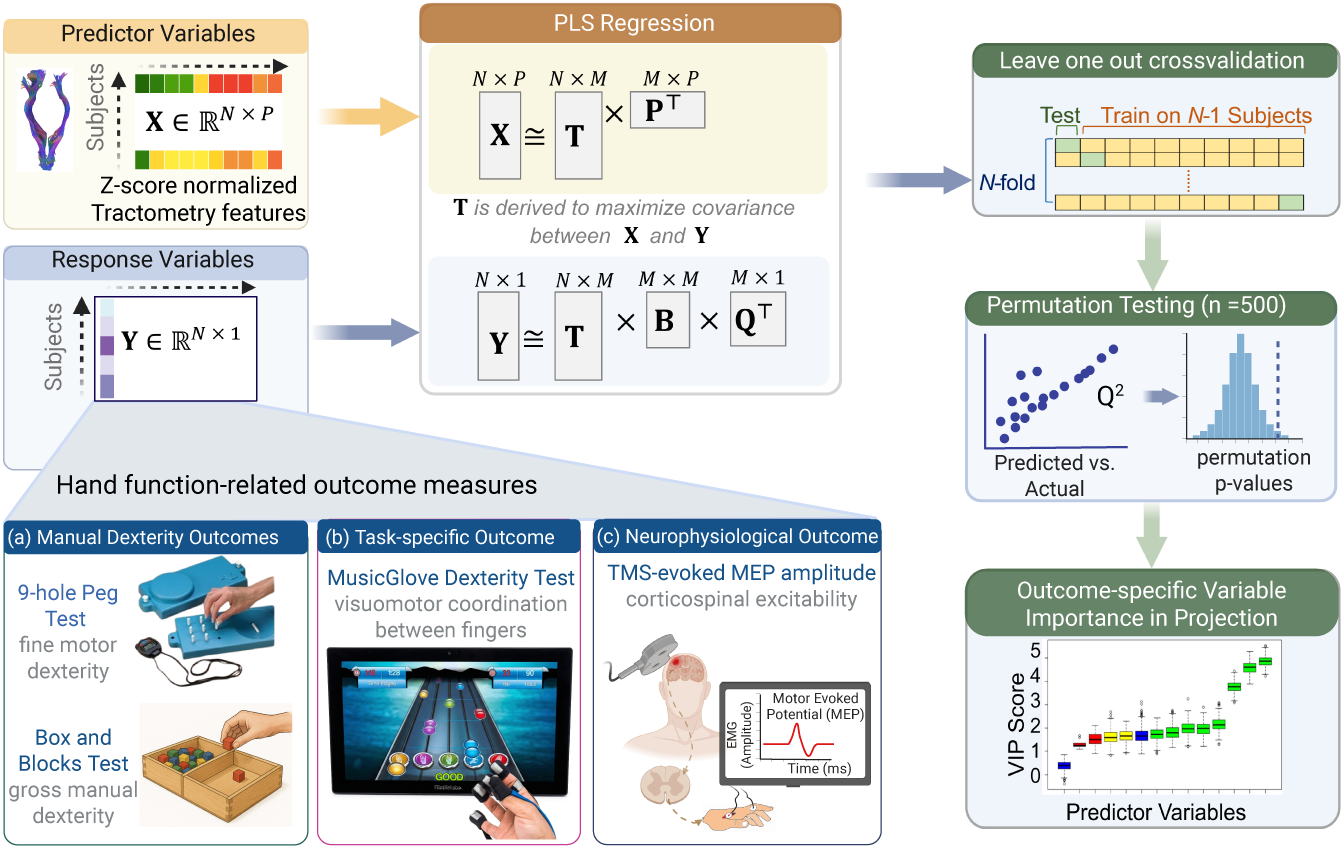
Overview of the partial least squares regression (PLSR) analytical workflow used to relate structural tractometry features to hand function outcomes. Z-score normalized tract-level diffusion and tractometry metrics (**X**) were used to predict individual outcome measures (**Y**), including manual dexterity (9-Hole Peg Test, Box and Block Test), task-specific performance (MusicGlove Dexterity Test), and neurophysiological function (TMS-evoked MEP amplitude). PLSR decomposes the predictor and response matrices into latent components that maximize covariance between **X** and **Y**. In the matrix notation, *N* denotes participants, *p* denotes the number of predictors, and *M* denotes the number of retained latent components; **T** denotes the latent score matrix, **P** the predictor-loading matrix, **B** the regression or inner-relation matrix, and **Q** the response-loading matrix. Model performance was evaluated using leave-one-out cross-validation (*Q*^2^) and permutation testing (*n* = 500). Outcome-specific variable importance in projection (VIP) scores were computed to identify the tractometry features contributing most strongly to each model. Figure created with BioRender.com.

Models were implemented in R using the pls package [50]. Predictive performance was estimated using nested leave-one-out cross-validation (LOOCV). In each outer fold, one participant was held out for testing, while the remaining participants were used to select the number of latent components and refit the model. Within each fold, predictors were z-score normalized before model fitting, and no data-driven feature selection was performed before cross-validation. Held-out predictions were then aggregated across folds to compute cross-validated model performance.

Model performance was summarized using cross-validated *Q*^2^, RMSE, MAE, the observed–predicted correlation coefficient (*r*), and the number of retained latent components. Values of *Q*^2^ *>* 0.1 were interpreted as evidence of meaningful internal predictive relevance [51], but because of the small sample size, positive *Q*^2^ values were considered hypothesis-generating rather than externally validated prediction. Statistical significance was assessed using 500 outcome-label permutations in which the full nested cross-validation procedure was repeated. Predictor importance was summarized using VIP scores, and bootstrap resampling was used to describe the stability of VIP rankings [52]. No inferential testing was performed on individual predictors. Additional implementation details, including the *Q*^2^ formula, component-selection procedure, permutation testing, and VIP bootstrap procedure, are provided in Supplementary Materials.

### 2.6 Correlational Tractography

As exploratory analyses, diffusion MRI connectometry was performed using DSI Studio (Chen release 2024) following the framework described by Yeh et al. [53], to identify WM segments in which local QA was associated with dexterity performance. BBT and MGDT were examined independently within the stroke and TBI cohorts. Associations were evaluated using nonparametric Spearman partial correlations, controlling for age and time since injury via multiple regression. The covariates were selected a priori because the cohorts differed in age and chronicity; however, because the within-cohort sample sizes were small, covariate-adjusted connectometry findings were interpreted cautiously.

A T-score threshold of 2.5 was applied to initiate tracking, and deterministic fiber tracking [14] was used to reconstruct correlational tracts. Tracks were filtered using topology-informed pruning (16 iterations) [41], with a length threshold of 30 voxels. To estimate statistical significance, 4000 permutations were performed to generate a null distribution of track length. The FDR threshold was set at 0.0125 to account for the four planned connectometry analyses (BBT and MGDT in the stroke and TBI cohorts). Results surviving this threshold were interpreted as statistically supported exploratory associations. Tract patterns that did not survive FDR correction were not used to support primary conclusions and were described only as descriptive observations when shown for completeness. Cerebellar regions were excluded from seeding to restrict analyses to supratentorial WM. For anatomical interpretation, significant tracts were classified using the HCP842 tractography atlas within DSI Studio, and percentage overlap with atlas-defined tracts was extracted for descriptive purposes.

## 3 Results

Participant demographic, clinical, behavioral, and neurophysiological characteristics are summarized in Table 1. The groups differed in age (stroke: 57.8 *±* 7.2 years; TBI: 46.8 *±* 11.2 years; *p* = 0.027) and time since injury (stroke: 25.0 *±* 12.4 months; TBI: 186.8 *±* 117.8 months; *p* = 0.003). Age and time since injury were therefore included as covariates in PLSR and connectometry analyses. Although the sex distribution comparison did not reach statistical significance (*p* = 0.082), the TBI cohort consisted exclusively of male participants, whereas the stroke cohort included both male and female participants. Handedness distribution did not differ between groups (*p* = 1.000). The TBI cohort showed higher MGDT scores before correction for multiple comparisons, whereas BBT, 9HPT, FDI MVC, MEP amplitude, and TMS motor threshold did not differ significantly between groups. After correction across behavioral and neurophysiological comparisons, no outcome measure showed a statistically significant between-group difference. Thus, the cohorts showed overlapping ranges of hand motor impairment but differed in demographic and injury-related characteristics. The corresponding distribution plots are provided in Supplementary Fig. S1.

### 3.1 Lesion Burden in Stroke

Lesion mapping was performed to describe the anatomical burden of stroke lesions and to help interpret the diffusion MRI findings. Lesion overlap demonstrated substantial interindividual heterogeneity, with the greatest overlap involving left frontal, insular, perisylvian, and subcortical regions (Fig. 4a). Projection of these lesion masks onto the HCP-1065 tractography atlas using LQT showed that focal lesions were associated with broader patterns of estimated WM disconnection extending beyond the visible lesion core (Fig. 4b). The group mean disconnection map most prominently involved projection and association pathways traversing perilesional regions, indicating that the structural burden of stroke was not confined to the lesioned gray matter territory alone. These maps are presented as descriptive anatomical context and were not used for formal lesion-symptom inference.

### 3.2 PLSR predictive modeling results

Across outcomes in the stroke cohort, PLSR showed positive internal cross-validated performance for the two primary behavioral dexterity measures but not for secondary neurophysiological or fine-motor outcomes (Table 2). BBT performance showed the strongest predictive relevance (*Q*^2^ = 0.69, *r* = 0.85, permutation *p* = 0.010). MGDT also showed positive predictive relevance, although the effect was more modest (*Q*^2^ = 0.22, *r* = 0.72, permutation *p* = 0.008). In contrast, models predicting 9HPT, FDI MVC, and MEP amplitude yielded negative *Q*^2^ values, indicating no predicitive relevance. These findings are illustrated in Fig. 3. For BBT, LOOCV-predicted scores tracked the observed scores and the observed *Q*^2^ exceeded the permutation-derived null distribution. VIP analysis of the full model indicated that q-space imaging metrics (QA and ISO) of the bilateral CST and tract-volume asymmetry contributed most strongly to prediction. A similar but weaker pattern was observed for MGDT, with bilateral CST isotropic diffusion and left CST QA ranking highest in variable importance. Bootstrap confidence intervals for the PLSR model performance are reported in Supplementary Table S2.

**Fig. 3.**
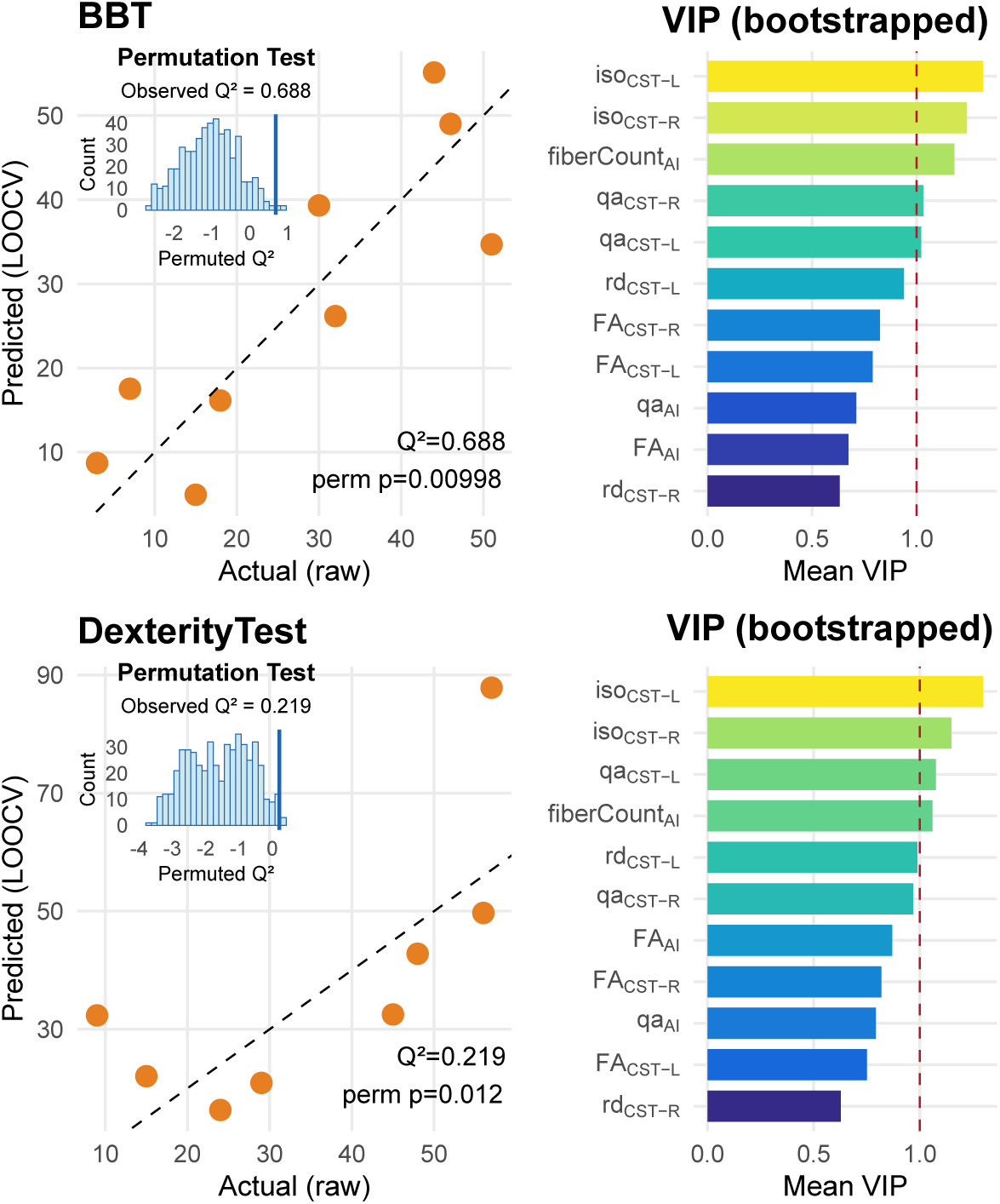
Observed versus LOOCV-predicted raw outcome scores for exploratory PLSR models with *Q*^2^ *>* 0.1 in the stroke cohort. Left panels show scatter plots of observed (x-axis) versus predicted (y-axis) raw scores for (top) BBT and (bottom) MGDT, with the dashed line indicating the identity line (perfect prediction), and each orange dot represents a subject. Insets display permutation distributions of *Q*^2^ (500 permutations), with the observed *Q*^2^ indicated; permutation *p*-values reflect the proportion of null models exceeding the observed *Q*^2^. Right panels show bootstrap-derived VIP scores for tract-specific predictors. Bars represent mean VIP values across bootstrap iterations, with the dashed vertical line indicating the threshold for influential predictors (VIP = 1). Predictors include diffusion metrics derived from bilateral CST, including FA, QA, RD, ISO, tract count, and asymmetry indices (AI) computed between contralesional and ipsilesional CST.

**Fig. 4.**
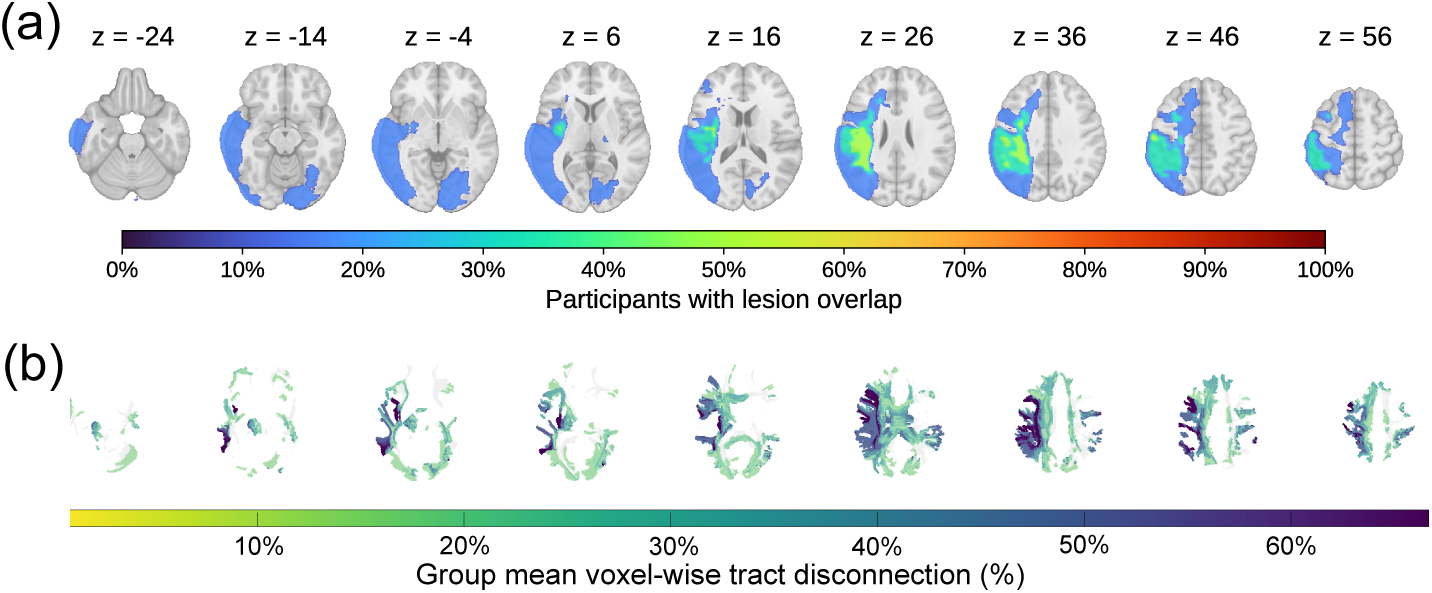
Group-level lesion topography and lesion-induced structural disconnection in the stroke cohort. (a) Lesion overlap map derived from automated segmentation of T1-weighted MRI scans and transformed to MNI152 standard space. Color intensity indicates the percentage of participants with lesion involvement at each voxel. (b) Group mean voxel-wise tract disconnection map generated using LQT by projecting subject-specific lesion masks onto the HCP-1065 population tractography atlas. Values indicate the mean percentage of streamlines estimated to intersect lesioned tissue across participants.

**Table 2.**
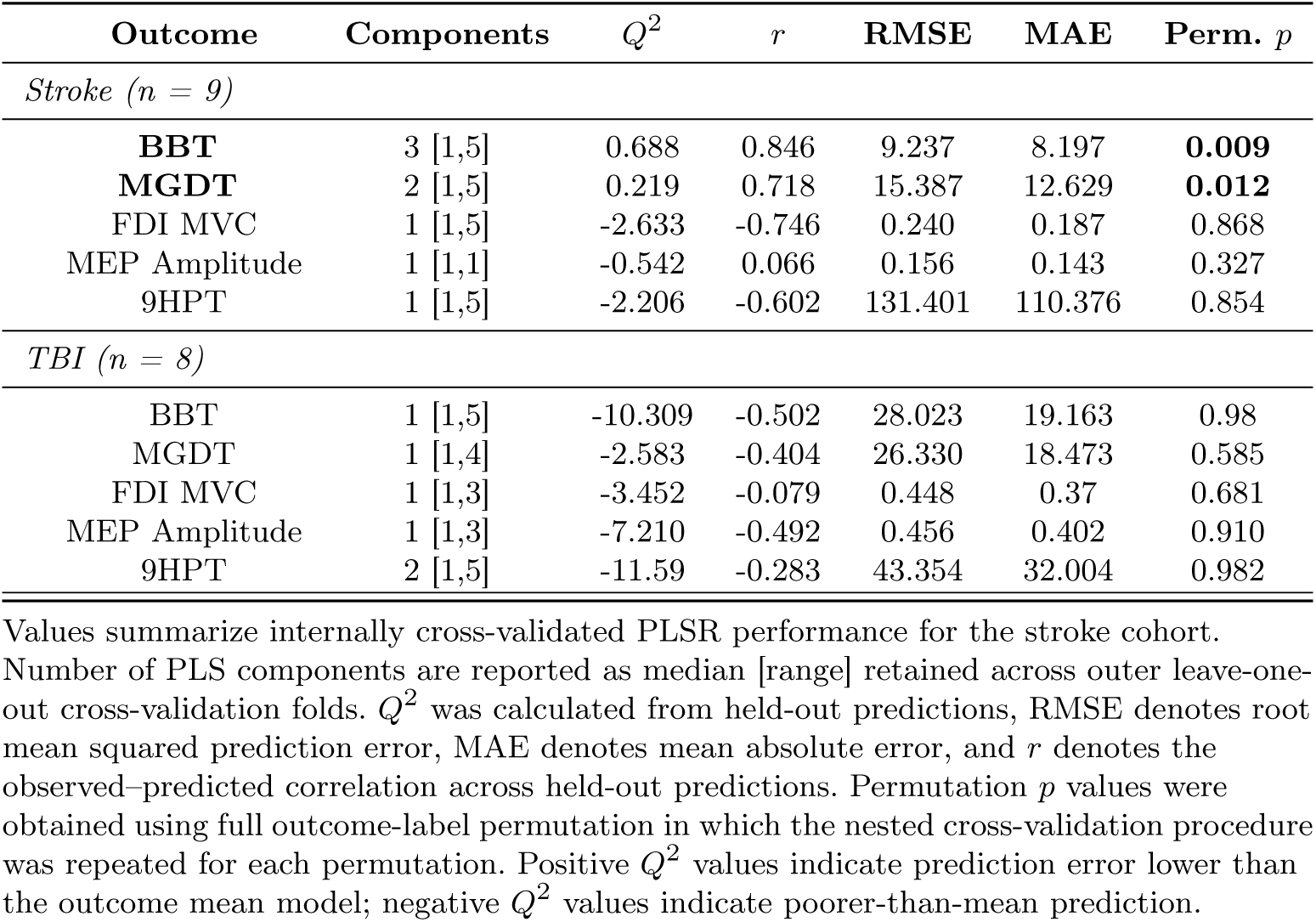
Leave-one-out cross-validated PLSR model performance in the stroke cohort.

In contrast to stroke, no outcome in the TBI cohort demonstrated positive predictive relevance (Table 2). All models yielded negative *Q*^2^ values, indicating that LOOCV prediction performed worse than mean-only estimation. Correlations between observed and predicted scores were weak to moderate and inconsistent in direction, and nonparametric permutation testing did not support statistical significance for any outcome. Although some diffusion metrics exhibited moderate VIP scores within fully fitted TBI models, these VIP patterns were not interpreted because the corresponding cross-validated models did not show predictive relevance. These results suggest that CST-restricted tractometry was insufficient to explain functional or neurophysiological variability in this small chronic TBI cohort.

### 3.3 Correlational Tractography

#### 3.3.1 Stroke Cohort

To investigate whether dexterity was associated with pathways beyond the CST, we conducted correlational tractography within the stroke cohort (Fig. 5). BBT scores were positively correlated with QA in interhemispheric and limbic-association pathways, most prominently the left cingulum frontal-parahippocampal tract, corpus callosum body, and corpus callosum tapetum, with additional contributions from the right inferior fronto-occipital fasciculus, left arcuate fasciculus, forceps major/minor, bilateral cingulum segments, and the right CST. Overall, BBT scores were associated with a distributed network centered on callosal and cingulo-association pathways, with smaller representation from long-range association and descending motor tracts.

**Fig. 5.**
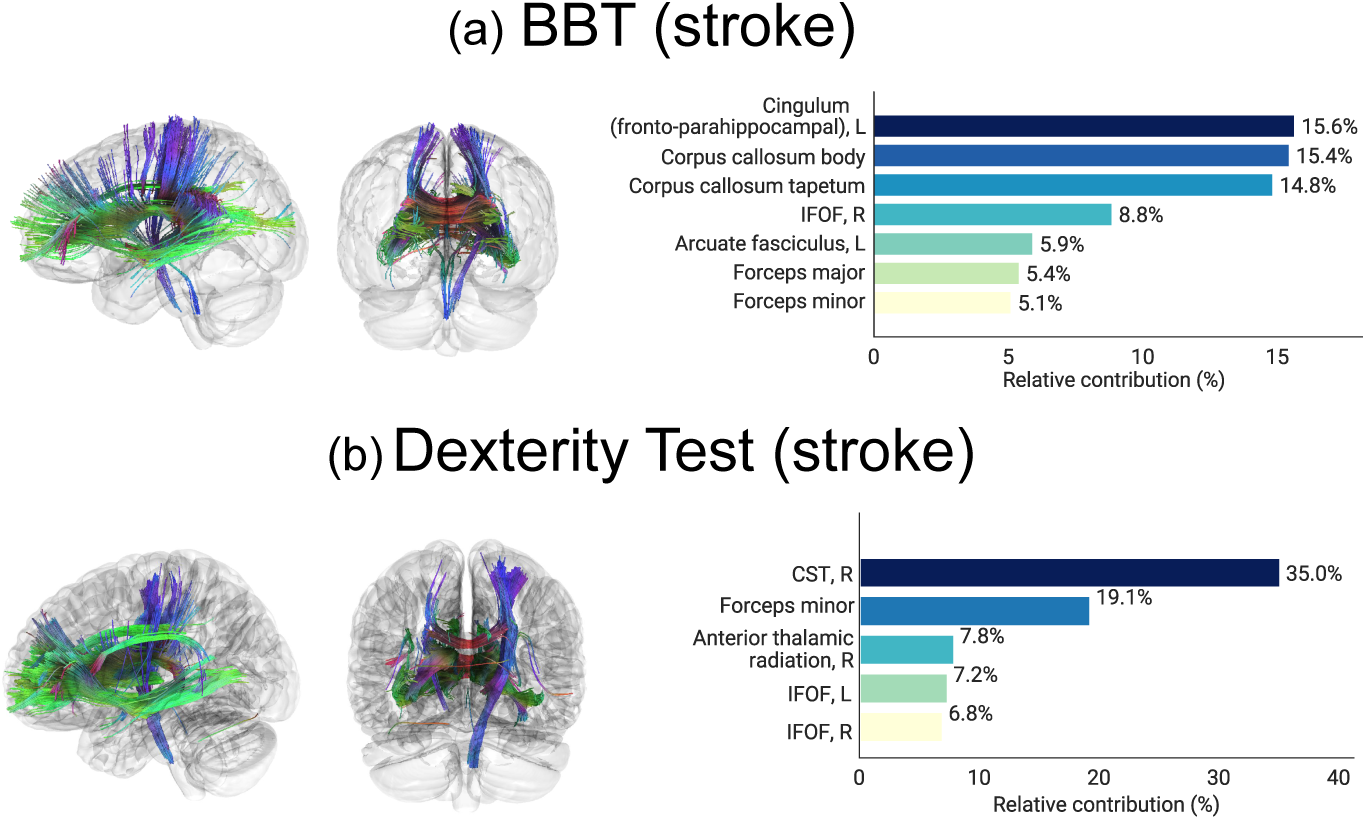
White matter pathways associated with dexterity in the stroke cohort, identified using exploratory correlational tractography. Significant tracts positively correlated with (a) BBT and (b) MGDT scores are shown in lateral and coronal views. Horizontal bar plots summarize the relative representation (%) of atlas-recognized tracts (*>* 5%) identified using the HCP842 atlas; percentages describe atlas overlap among significant streamlines and should not be interpreted as biological contribution or absolute tract strength. In stroke, dexterity-related findings involved prominent commissural pathways, including the corpus callosum body, tapetum, and forceps fibers, together with association and projection pathways such as the cingulum, arcuate fasciculus, inferior fronto-occipital fasciculus (IFOF), anterior thalamic radiation, and corticospinal tract (CST).

For MGDT, scores were positively correlated with QA in the right CST and forceps minor, with additional positively associated segments in the right anterior thalamic radiation, bilateral inferior fronto-occipital fasciculi, right parietal corticopontine tract, right reticular tract, and smaller contributions from cingulum, uncinate fasciculus, fornix, corticobulbar, and cerebellar-thalamic pathway segments. Compared with BBT, MGDT showed relatively greater representation of descending motor and right-hemispheric projection pathways based on the atlas-derived tract percentages summarized in Fig. 5. No significant negative correlations were identified in the stroke analyses. These findings indicate that, within the stroke cohort, BBT and MGDT were associated with partially overlapping but behavior-specific structural networks.

### 3.4 TBI Cohort

To examine whether distributed WM pathways were associated with dexterity performance in the TBI cohort, we performed correlational tractography (Fig. 6). Statistically supported tracts surviving FDR correction were identified for BBT but not MGDT. For BBT, higher scores were associated with greater QA in a distributed set of commissural and association pathways, including prominent contributions from the corpus callosum and right-lateralized frontoparietal association tracts. Segments of the superior longitudinal fasciculus and frontal aslant tract showed substantial representation, with additional involvement of cingulum subdivisions, fornix, and projection pathways including the dentatorubrothalamic tract. These results suggest that BBT performance in this small TBI cohort was associated with distributed commissural, association, limbic, and cerebellar-thalamic pathway segments rather than being dominated by CST contributions. No significant negative correlations were identified. The MGDT connectometry analysis in TBI did not survive the prespecified FDR threshold. Therefore, the MGDT tract pattern shown in Fig. 6 is retained only as a descriptive visualization and is not used to support the primary TBI conclusions.

**Fig. 6.**
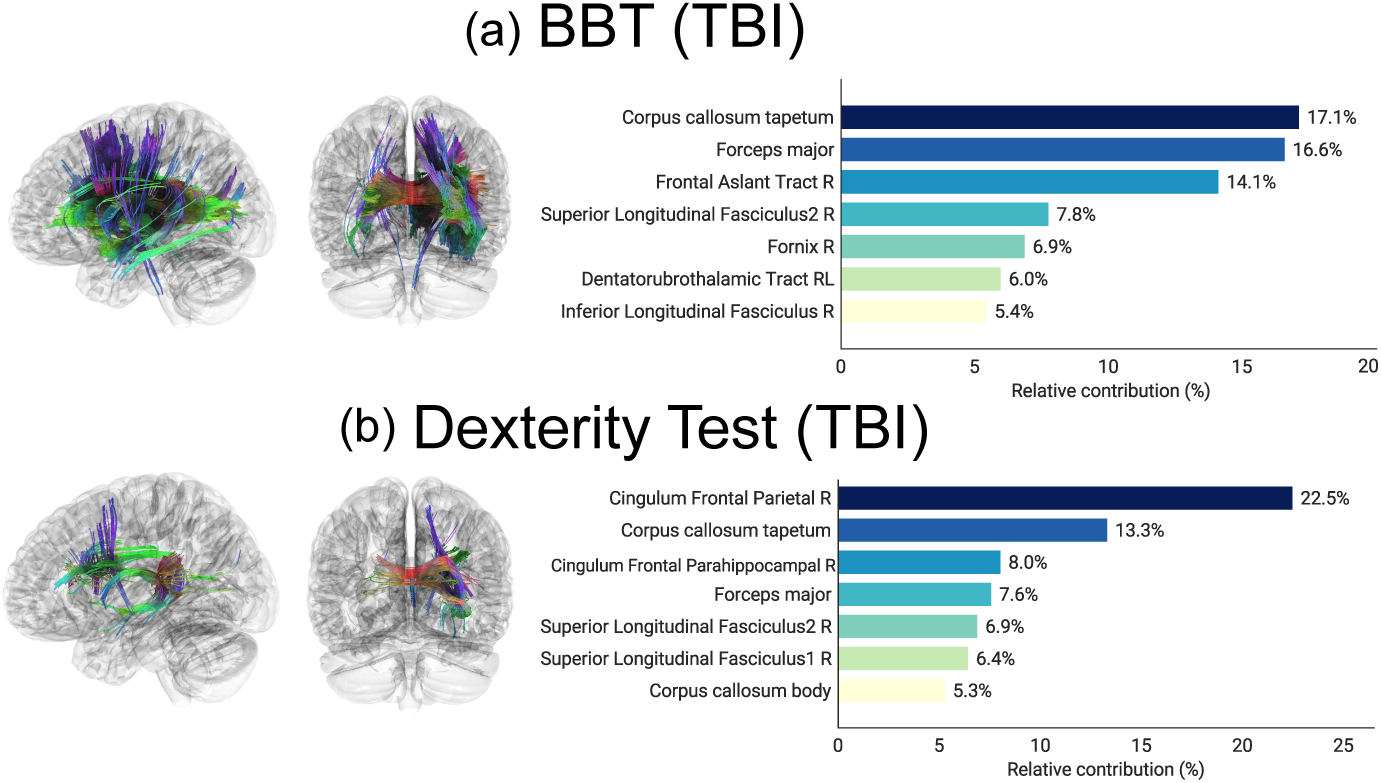
White matter pathways associated with dexterity in the TBI cohort, identified using correlational tractography. Statistically significant and FDR-corrected streamlines positively correlated with (a) BBT are shown in lateral and coronal views; panel (b) shows the MGDT analysis for completeness, because this analysis did not survive the prespecified FDR threshold. Horizontal bar plots summarize the relative representation of tracts (*>* 5%) recognized by the HCP842 atlas. In TBI, dexterity-related findings involved commissural pathways together with right-lateralized association and limbic tracts, including the frontal aslant tract, superior longitudinal fasciculus, cingulum sub- divisions, fornix, and dentatorubrothalamic tract.

## 4 Discussion

### 4.1 Predictive modeling of hand dexterity using CST tractometry

The main finding of this retrospective exploratory study is that CST-derived diffusion and tractometry features showed internally cross-validated predictive relevance for two functional dexterity measures in the chronic stroke cohort, particularly BBT, whereas CST-specific models did not predict behavioral or neurophysiological outcomes in the chronic TBI cohort. This main finding support the view that residual hand dexterity after acquired brain injury is not fully captured by a single tract or single behavioral measure.

The positive stroke PLSR findings are concordant with prior studies linking CST integrity to residual upper-limb function and recovery potential [19, 42, 54]. The current results extend this literature by showing that multivariate CST-derived features retained positive predictive relevance under nested LOOCV rather than only showing in-sample associations. The contribution of asymmetry indices further suggests that bilateral imbalance in descending motor system integrity may carry information beyond any single unilateral metric, consistent with evidence that chronic stroke performance reflects the broader bilateral motor system rather than the ipsilesional CST alone [54, 55]. However, we acknowledge that the model estimates may be sensitive to individual participants and require external validation before being considered prognostic biomarkers.

Furthermore, the dissociation between behavioral and neurophysiological outcomes is also informative. Although CST metrics showed predictive relevance for functional dexterity, they did not predict MEP amplitude, FDI MVC, or 9HPT completion time. TMS-derived physiology depends not only on tract microstructure but also on corticospinal excitability, synaptic efficacy, lesion proximity to M1, medication effects, stimulation geometry, and peripheral muscle factors. Preserved WM integrity may therefore support descending transmission without fully explaining variability in TMS or EMG outcomes. Measurement variability in neurophysiological outcomes may also have further contributed to these null findings.

The absence of predictive relevance in TBI should be interpreted cautiously. Unlike stroke, TBI often produces diffuse and spatially heterogeneous WM disruption beyond conventional motor pathways. Under these conditions, a CST-specific model may not be informative enough in TBI, in which chronic deficits can arise from distributed disconnection rather than focal injury to a single tract [56, 57]. Our null TBI PLSR findings are best interpreted as evidence that CST tractometry was not sufficient in this small sample, which was exclusively male, and much more chronic than the stroke cohort.

### 4.2 Network-level patterns of correlational tractography

Connectometry complemented the PLSR results by showing that dexterity was associated with distributed WM segments beyond the CST. In stroke, higher BBT scores were associated mainly with greater QA in callosal and cingulum-related pathways, including the corpus callosum body, tapetum, forceps fibers, and cingulum segments. This pattern suggests that BBT performance reflects not only CST microstructure but also networks supporting interhemispheric communication, visuomotor integration, sequencing, and higher-order motor planning. This interpretation is consistent with evidence that corpus callosum integrity contributes to arm function and bimanual performance after stroke [55, 58, 59], and with network-based work showing that post-stroke behavior can depend on preserved interhemispheric and association connectivity beyond lesion topography alone [8, 29].

The stroke MGDT findings showed a greater representation of the right CST, forceps minor, anterior thalamic radiation, and bilateral inferior fronto-occipital fasciculi. We postulate that this divergence between the BBT and MGDT-specific patterns could be attributed to the greater demands on rapid sensorimotor execution, finger sequencing, and sustained descending motor control during the MGDT task while still requiring visuomotor and executive-control pathways. This task dependence supports behavior-specific structure-function relation and aligns with evidence that broader fronto-parietal systems can support motor performance when CST integrity is compromised [8].

In TBI, statistically supported connectometry findings were limited to BBT. Higher BBT scores were associated with distributed commissural, association, limbic, and cerebellar-thalamic pathway segments, including corpus callosum, frontal aslant tract, superior longitudinal fasciculus, cingulum subdivisions, fornix, anterior commissure, and dentatorubrothalamic pathways. This pattern is broadly concordant with TBI as a disorder of large-scale network disconnection [56, 57]. Prior studies have linked corpus callosum microstructure to bimanual impairment after TBI and suggested that motor recovery after diffuse axonal injury may rely on widespread reorganization rather than a single descending tract [5, 26, 60]. However, the MGDT connectometry analysis in TBI did not survive FDR correction and should not be used to support conclusions about TBI dexterity networks.

### 4.3 Lesion Burden and Topography

The lesion analysis provides anatomical context for linking the focal CST-based PLSR findings with the broader connectometry results. Lesion network mapping studies emphasize that focal stroke lesions can disrupt structurally connected regions and WM pathways beyond the visible lesion core [43, 48, 49]. Consistent with this view, projecting individual lesion masks onto a population tractography atlas showed that spatially heterogeneous stroke lesions produced broader estimated disconnection across projection and association pathways. This supports the interpretation that poststroke dexterity impairment is not fully captured by lesion location or CST tractometry alone, but is constrained by the broader residual WM network.

These lesion-disconnectivity maps should be interpreted as anatomical context rather than as a direct replication of the connectometry findings. LQT estimates which pathways are likely interrupted by lesions using a normative tractography atlas, whereas connectometry identifies segments in participants’ own diffusion data where preserved local QA correlates with hand function. The partial overlap between lesion burden in projection pathways and CST-based PLSR findings is consistent with prior evidence linking CST lesion load to poststroke motor outcome [49, 61]. Likewise, the broader disconnection pattern supports a network-level interpretation of poststroke motor impairment, consistent with recent work emphasizing distributed ipsilesional and interhemispheric connectivity in recovery [29, 30]. Thus, lesion-disconnectivity analysis strengthens the anatomical interpretation of the stroke findings, while remaining secondary and descriptive analysis.

### 4.4 Limitations and Future Directions

Several limitations should be considered. First, the sample size was small, and the analyses were retrospective. The resulting estimates are likely sensitive to individual participants, and the models should be viewed as hypothesis-generating rather than confirmatory. Second, the stroke and TBI cohorts were not matched for age, sex, or time since injury; the TBI cohort was exclusively male and substantially more chronic. These factors necessitate cautious interpretation of cohort-specific patterns. Third, although age and time since injury were included as covariates in connectometry, covariate adjustment in small samples cannot fully remove confounding. Accordingly, the generalizability of these findings is limited by the single-center design, small sample size, chronic-stage participants, and demographic imbalance between cohorts. The present findings should not be generalized to acute or subacute injury, more severe impairment, demographically balanced populations, or clinical prognostic settings without replication in larger prospective cohorts.

Fourth, the PLSR models were internally cross-validated but not externally validated. While LOOCV is useful in small samples, it can still have high variance and should not be used for clinical prognosis. Fifth, lesion-disconnectivity maps were available only for the stroke cohort and were used descriptively rather than as predictors. Finally, tractography-derived measures such as streamline count, tract volume, and tract length are reconstruction-dependent and should not be interpreted as direct anatomical measures of axonal number.

Despite these limitations, the study provides a useful framework for future work. In larger prospective cohorts, CST tractometry, lesion-disconnectivity metrics, whole-brain connectometry, and graph-theoretic network measures should be evaluated together, with prespecified primary outcomes, external validation, and sensitivity analyses for age, sex, chronicity, lesion burden, and injury severity. Such work will be necessary before diffusion MRI markers can be used for individualized prognosis or treatment stratification in neurorehabilitation.

## 5 Conclusion

In this small retrospective baseline study, CST-derived diffusion and tractometry features showed internally cross-validated predictive relevance for functional hand dexterity in chronic stroke, particularly BBT and MGDT, whereas CST-restricted models did not predict outcomes in the TBI cohort. Whole-brain connectometry suggested that dexterity was also associated with distributed callosal, association, and projection pathways, with behavior-specific patterns across BBT and MGDT in stroke and statistically supported distributed BBT associations in TBI; whereas the MGDT analysis was null after correction in TBI. These findings support the value of combining focal CST tractometry with whole-brain network analyses, but they should be interpreted as preliminary and exploratory. Larger prospective studies with external validation are needed to determine whether these candidate diffusion MRI markers can improve prognosis or guide individualized neurorehabilitation strategies.

## Acknowledgments

The authors thank the participants for their involvement in the study procedures and acknowledge support from the study teams at Kessler Foundation and the Rocco Ortenzio Neuroimaging Center.

## Reporting guideline

This retrospective observational analysis was reported in accordance with STROBE principles where applicable. A completed checklist is included as supplementary material.

## Data availability statement

Raw MRI data and participant medical history are not publicly available because of IRB restrictions and participant privacy considerations. Deidentified derived data and analysis scripts may be made available from the corresponding author upon reasonable request, subject to institutional approval and data-use agreement.

